# Adenosine metabolized from extracellular ATP promotes type 2 immunity through triggering A_2B_AR signaling on intestinal epithelial cells

**DOI:** 10.1101/2021.01.24.428000

**Authors:** Darine W. El-Naccache, Fei Chen, Mark Palma, Alexander Lemenze, Wenhui Wu, Pankaj K. Mishra, Holger K. Eltzschig, Simon C. Robson, Francesco Di Virgilio, György Haskó, William C. Gause

## Abstract

Multicellular intestinal nematode parasites can cross the epithelial barrier potentially causing tissue damage and release of danger associated molecular patterns (DAMPs) that may promote type 2 responses and host protective immunity. We investigated whether adenosine specifically binding the A_2B_ adenosine receptor (A_2B_AR) on epithelial cells played an important role in driving intestinal immunity. Specific blockade of epithelial cell A_2B_AR inhibited the host protective memory response to the enteric helminth, *Heligmosomoides polygyrus bakeri*, including disruption of granuloma development at the host:parasite interface during the transient tissue dwelling larval stage. Memory T cell development was blocked during the primary response and transcriptional analyses revealed profound impairment of A_2B_AR signaling in epithelial cells and reduced type 2 markers by 24 hours after inoculation. Extracellular ATP was visualized by 24 hours after inoculation and shown in CD39 deficient mice to be critical for the adenosine production mediating initiation of type 2 immunity.

## Introduction

Soil-transmitted helminth infections are a global health issue resulting in high rates of morbidity with an estimated 1.5 billion people infected worldwide. Infected individuals commonly suffer from malnutrition, anemia and severe immunopathology. Indeed, helminth infections remain a significant concern for human and livestock health. The incidence of high reinfection rates and increased susceptibility to co-infections with other infectious agents warrants the need for the development of novel improved treatments and prevention strategies. Helminth infections and other insults such as allergens and sterile particles can induce a type 2 immune response in both humans and experimental mouse models. This response is characterized by upregulation of interleukin (IL)-4, IL-5, and IL-13 by lymphocytes and myeloid cells, alternatively activated (M2) macrophages, and fibrosis [1, 2].

Recent studies have revealed multiple essential mechanisms contributing to the initiation and development of the host protective type 2 immune response to enteric helminths. Specialized intestinal epithelial tuft cells can apparently sense invading parasites in the lumen. This triggers tuft cell secretion of cysteinyl leukotrienes, which in conjunction with their constitutive secretion of IL-25 can promote IL-13 production by activated ILC2s, leading to tuft cell hyperplasia [3–5]. As these large multicellular parasites rupture and migrate through the intestinal tissue, they can cause tissue damage and release of danger associated molecular patterns (DAMPs), which can trigger immune cell activation. Recent studies have suggested that DAMPs, such as extracellular ATP (eATP) [6], can bind their receptors to trigger an immune response. eATP, released from stressed/damaged cells, binds the ATP-P2X7 receptor on mast cells inducing IL-33 release. IL-33 likely triggered through a number of mechanisms can then act on a variety of innate immune cells including type 2 innate lymphoid cells (ILC2s) and myeloid cells leading to production of type 2 cytokines and the development of type 2 immune responses. Trefoil factors 2 and 3 can also function as DAMPs playing an essential role in driving helminth induced type 2 immune responses[7, 8].

Adenosine is a purine nucleoside that contributes to numerous modulatory functions and can regulate various biological processes by binding to the G-protein coupled cell surface adenosine receptors (A_1_, A_2A_, A_2B_ and A_3_ –AR). Extracellular adenosine levels increase during tissue injury, inflammatory stress, and mechanical insults. The accumulation of extracellular adenosine can be due to its release from cells or the release of ATP followed by its catabolism to adenosine through cell surface ectonucleotidases, CD39 and CD73 [9]. Recent studies have demonstrated a potential role for the A_2B_ adenosine receptor in modulating the type 2 immune response. *In vitro* studies have demonstrated that adenosine can promote the development of alternatively activated (M2) macrophages by binding to A_2B_AR expressed on the macrophage [10]. A_2B_AR signaling on mast cells has also been shown induce their production of IL-4 *in vitro* [11]. We have previously reported that adenosine can act as a DAMP through the A_2B_AR to initiate type 2 immune responses. Our studies demonstrated that A_2B_AR deficient mice (A_2B_AR^−/−^) have an impaired type 2 immune response and delayed helminth expulsion [9]. However, the cell types expressing A_2B_AR receptors that participate in this initiation and how the adenosine-A_2B_AR axis drives memory type 2 immune responses in helminth infections remains unknown.

The natural murine gastrointestinal nematode parasite *Heligmosomoides polygyrus bakeri (Hpb)* is an established model to study the type 2 immune response and has been an important experimental for studying both memory and initiation of type 2 immunity. After peroral inoculation, L3 parasites quickly invade intestinal tissues migrating to the submucosa where they develop for 8 days, when they return as adults to the intestinal lumen. Here they mate and produce eggs, which are passed in the feces, and can reproduce and persist in the lumen for prolonged periods [12]. Secondary inoculation after drug-induced worm clearance triggers a potent memory CD4 T cell-dependent type 2 response that mediates effective worm clearance by day 14 after inoculation. During the initial tissue dwelling phase, macrophages quickly surround the developing larvae after secondary inoculation forming a type 2 granuloma which contributes to clearance [13–15].

We now report that A_2B_AR expression by epithelial cells, but not macrophages, is essential for the development of an effective memory response after *Hpb* inoculation. We further show that A_2B_AR epithelial cell signaling mediates the development of memory T cells during the primary response but is not required for subsequent memory T cell activation after secondary inoculation. During the primary response, type 2 immunity was dependent on epithelial A_2B_AR signaling as early as 24 hours after inoculation and was associated with marked increases in eATP production. Blockade of CD39 similarly inhibited memory type 2 immune responses, thus demonstrating the importance of eATP as a catabolic source of extracellular adenosine.

## Results

### Intestinal epithelial A_2B_AR signaling promotes host protective memory immune response to intestinal helminth Hpb

*Hpb* is a strictly enteric murine intestinal nematode parasite that triggers a potent and polarized type 2 immune response. After peroral inoculation, larval parasites invade intestinal tissues residing in the submucosa for 8 days, and then return as adults to the intestinal lumen where they persist for prolonged periods, matinag and producing eggs passed in the feces. Secondary inoculation after drug-induced worm clearance triggers a potent memory CD4 T cell-dependent type 2 response that mediates effective worm clearance [13, 14]. We have previously reported that mice deficient in adenosine A_2B_ receptor (A_2B_AR) have an impaired type 2 immune response and delayed worm expulsion following secondary inoculation [9]. Intestinal epithelial cells (IECs) [16–18] and innate immune cells, in particular macrophages [19], express A_2B_AR. Previous studies have suggested that A_2B_AR signaling drives differentiation of alternatively activated (M2) macrophages [10, 20, 21], which in turn mediate helminth expulsion during a memory type 2 immune response [14]. To examine whether A_2B_AR signaling in macrophages or epithelial cells played a significant role in the host protective memory response to *Hpb*, mice genetically deficient in A_2B_AR on intestinal epithelial cells (Villin^Cre^-A_2B_AR^fl/fl^) and mice with a deficiency in A_2B_AR in myeloid cells (LysM^Cre^ – A_2B_AR^fl/fl^) were infected with *Hpb*. Specifically, mice were inoculated orally with 200 L3 *Hpb* (1‘*Hpb*) and 14 days post infection, mice were administered the anti-helminthic drug pyrantel pamoate via oral gavage for 2 consecutive days, effectively removing all parasites from the gut, as described previously [14]. At six weeks post clearance, A_2B_AR lineage specific KO and control lineage specific Cre mice were challenged with a secondary inoculation of 200 L3 (2‘ *Hpb*). At this same time point, control groups of naïve A_2B_AR lineage specific and control Cre mice were given a primary *Hpb* inoculation. Fourteen days later all groups were analyzed for worm burden and effects on parasite metabolic activity and growth. As shown in Figure 1a, Villin^Cre^ control mice showed markedly reduced worm burdens at day 14 after secondary inoculation, consistent with an effective memory type 2 response. In contrast, Villin^Cre^-A_2B_AR^fl/fl^ mice administered a secondary inoculation showed significant increases in worm burden, not significantly different to Villin^Cre^-A_2B_AR^fl/fl^ given a primary inoculation only, indicating that the memory response was compromised. Although parasites are largely expulsed by day 14 in WT mice after a secondary inoculation, at day 11 they have only recently migrated back to the lumen after developing into adult worms during their tissue dwelling phase in the submucosa, which lasts up to day 8, and are thus readily detectable. In a separate experiment involving the same treatment groups as in Figure 1a, parasites were collected from the small intestine at day 11 after inoculation. Five adult worms in each treatment group were analyzed for overall metabolic activity by assessing ATP concentration levels, as previously described [22]. As shown in Figure 1b, ATP concentrations (nM) were significantly reduced in Villin^Cre^ mice after secondary inoculation, compared to Villin^Cre^ mice receiving only a primary inoculation. In marked contrast, parasite ATP concentration levels were not reduced in Villin^Cre^-A_2B_AR^fl/fl^ mice after secondary inoculation (compared to primary inoculation) and were significantly increased over Villin^Cre^ mice administered a secondary *Hpb* inoculation. Similar effects were observed when total worm protein was assessed (Fig. 1c). LysM^Cre^-A_2B_AR^fl/fl^ mice, where A_2B_AR is deleted in myeloid cells, were similarly assessed for host protective responses following *Hpb* inoculation. As show in Fig. 1d, worm burden was markedly reduced in both LysM^Cre^ and LysM^Cre^-A_2B_AR^fl/fl^ mice after secondary inoculation (compared to primary inoculation) as were the fitness of the worms assessed by ATP concentration levels (Fig. 1e) suggesting a strong polarized Th2 response. To assess whether type 2 responses were generally refractory to deletion of A_2B_ARs on macrophages, effects on a previously established peritoneal type 2 immune response to sterile inert microparticles (MPs) [23] were examined. Mice were inoculated in the peritoneal cavity with MPs, triggering myeloid cell infiltration and potent elevations in type 2 cytokines, as previously described [23]. As shown in supplementary Fig.1, at 48 hrs after inoculation pronounced elevations in type 2 cytokines and increases in M2 macrophages were blocked in MP-inoculated LysM^Cre^-A_2B_AR^fl/fl^ mice compared to inoculated LysM^Cre^ mice in this experimental model of sterile inflammation. Taken together, our data thus demonstrate a key role for intestinal epithelial cell-specific A_2B_AR signaling in the protective type 2 memory immune response to helminth infection.

**Figure 1.**
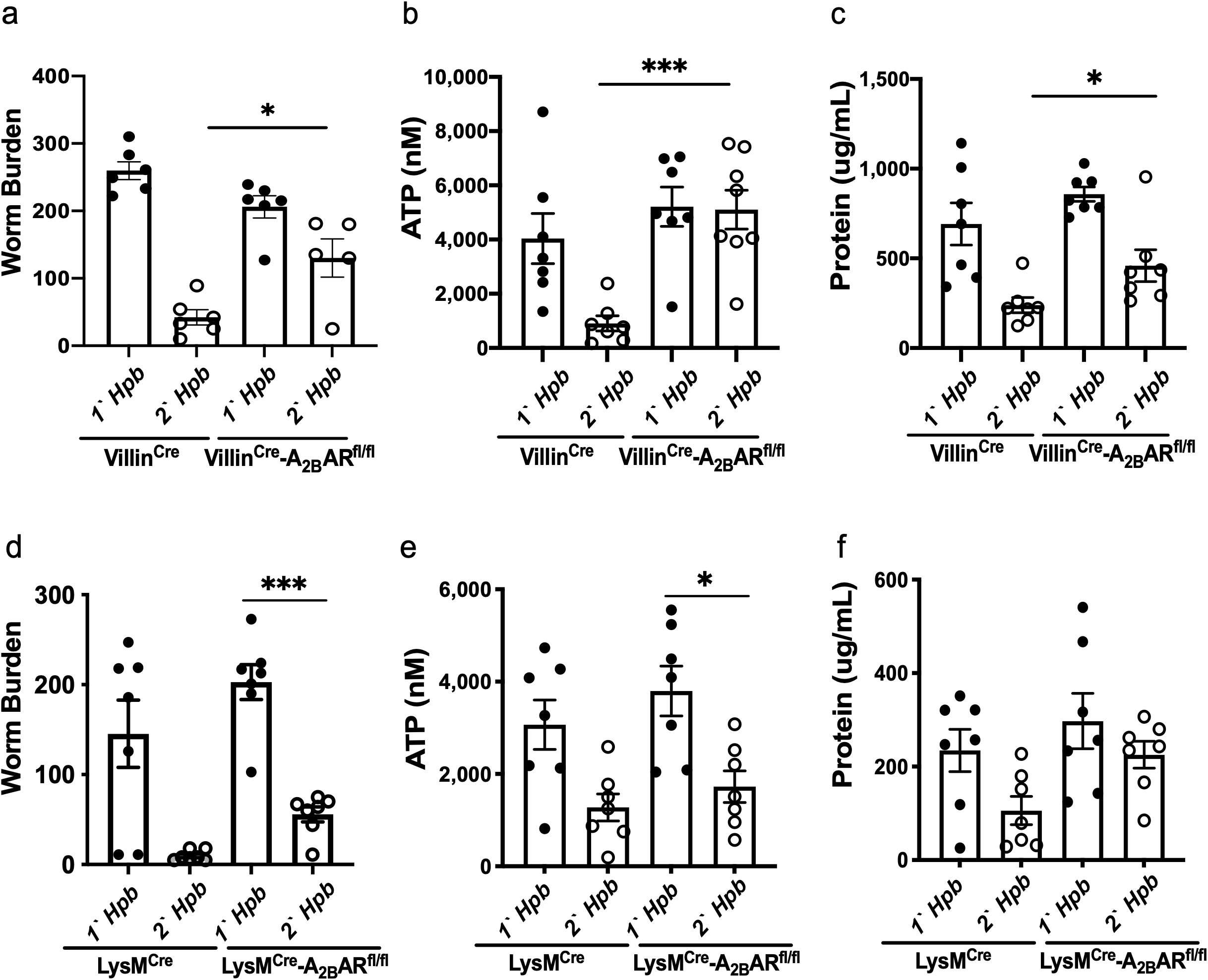
Intestinal epithelial A_2B_AR signaling promotes worm expulsion. (a-c) Villin^Cre^-A_2B_AR^fl/fl^ and corresponding controls Villin^Cre^ mice were orally inoculated with 200 L3 *Hpb* larvae, 14 days later, mice were treated with anti-helminthic drug, pyrantel pamoate, to expulse the parasites. At 6 weeks post clearance, mice were challenged with a secondary (2‘) *Hpb* inoculation and controls included mice given primary (1‘) *Hpb* inoculation. Resistance associated with memory response was assessed at day 14. Luminal worm burden was enumerated (a). Villin^Cre^-A_2B_AR^fl/fl^ and corresponding controls Villin^Cre^ mice were given a 1‘ and 2‘ *Hpb* inoculation as just described and the effects on parasite metabolic activity and growth were assessed at day 11. Individual parasites were isolated, female parasites were recovered from the lumen. ATP levels (nM) (b) and protein levels (c) were determined for five female worms per mouse. (d-f) LysM^cre^-A_2B_AR^fl/fl^ mice and corresponding controls LysM^Cre^ mice were orally inoculated with 200 L3 *Hpb* larvae, 14 days later, mice were treated with anti-helminthic drug, pyrantel pamoate for 2 constitutive days to expulse the parasites. At 6 weeks post clearance, mice were challenged with a secondary (2‘) *Hpb* inoculation and controls included mice given primary (1‘) *Hpb* inoculation. Resistance was assessed at day 14, luminal worm burden was enumerated (d). LysM^Cre^-A_2B_AR^fl/fl^ and corresponding controls LysM^Cre^ mice were given a 1‘ and 2‘ *Hpb* inoculation as just described and the effects on parasite metabolic activity and growth was assessed at day 11. Individual parasites were isolated, female parasites were recovered from the lumen. ATP levels (nM) (e) and protein levels (f) were determined for five female worms per mouse. Data shown are the mean and SEM from at least six individual mice per group (One-way ANOVA p<0.01, *p<0.05, **p<0.01, ***p<0.001, ****p<0.0001). Experiments were repeated at least two times with similar results.

### Intestinal epithelial A_2B_AR signaling promotes protective memory type 2 immune response at the host:parasite interface

CD4^+^ T cells and M2 macrophages rapidly accumulate at the host:parasite interface after secondary inoculation resulting in a distinct granulomatous structure surrounding the tissue-dwelling larvae by day 4 after secondary inoculation [14, 24]. To examine whether CD4 T cell-dependent granulomatous development occurring during the memory type 2 response was impacted by A_2B_AR deficiency in IECs, Villin^Cre^-A_2B_AR^fl/fl^ ad VillinCre control mice were administered a primary and secondary inoculation of *Hpb*, as described above. At day 4 after secondary inoculation, small intestines were collected and swiss roll cryosections were stained for F4/80 and CD206 to detect M2 macrophages. As shown in Figure 2a, immunofluorescent imaging of the granuloma showed dual-stained M2 macrophages already accumulating around the parasite in granulomas from VillinCre mice (Fig 2a). In marked contrast, in Villin^Cre^-A_2B_AR^fl/fl^ mice, although F4/80^+^ macrophages were readily detected, CD206 staining was markedly reduced, indicating an essential role for A_2B_AR signaling specifically in epithelial cells for the development of M2 macrophages within the granuloma. Punch biopsies of granulomas were collected as previously described and analyzed for mRNA expression of M2 markers and specific cytokines. Within the granuloma there was a significant decrease in M2 markers chi3I3, retnla, and Arg1 as well as a decrease in type 2 cytokines IL-4, IL-5 and IL-13 in Villin^Cre^-A_2B_AR^fl/fl^ compared to VillinCre controls (Fig. 2c-j).

**Figure 2.**
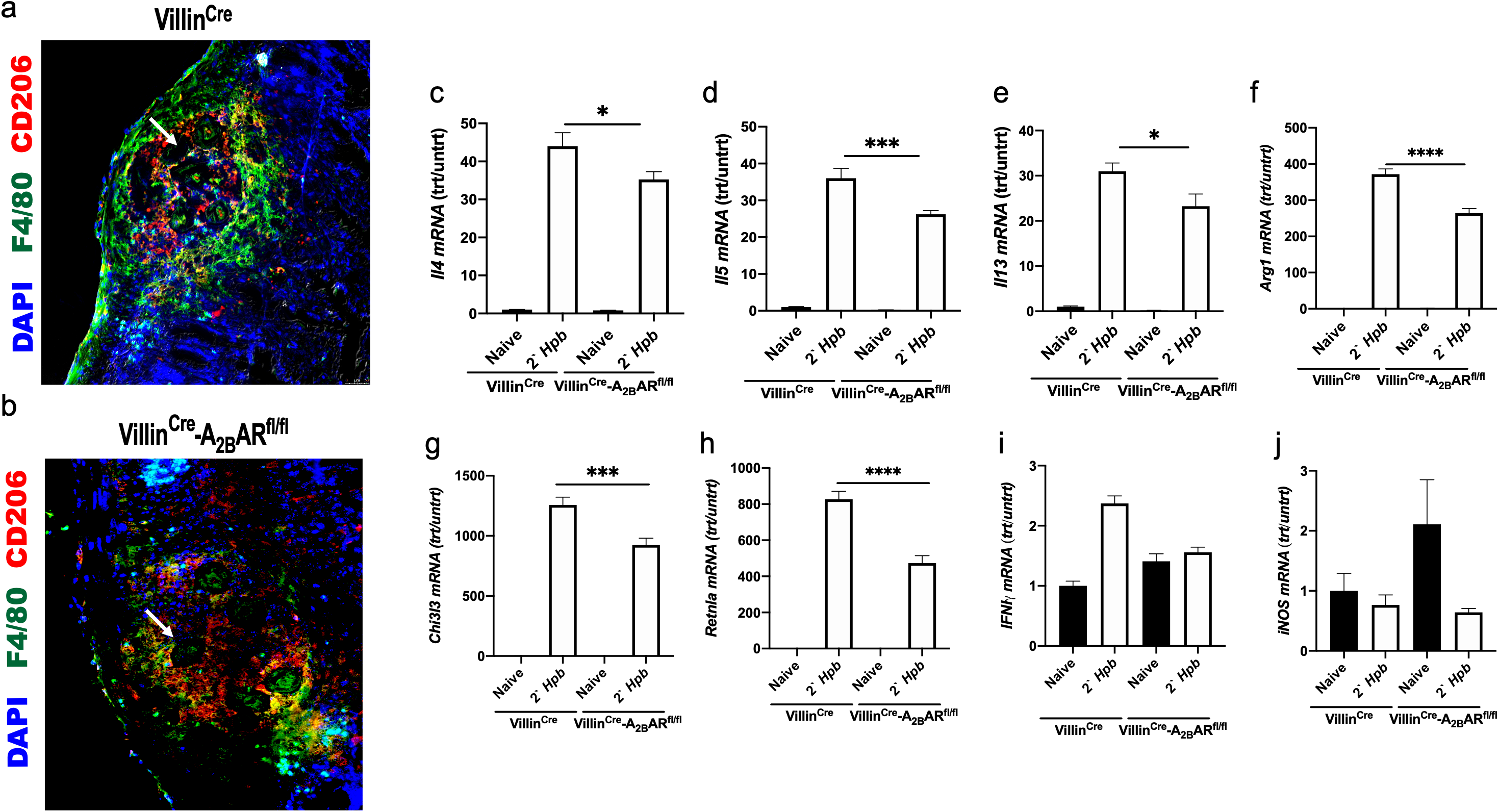
Intestinal epithelial A_2B_AR signaling promotes protective type 2 immune response at the host:parasite interface. Villin^Cre^-A_2B_AR^fl/fl^ and corresponding control Villin^Cre^ mice were orally inoculated with 200 L3 *Hpb* larvae and 14 days later, mice were treated with anti-helminthic drug, pyrantel pamoate, to expulse the parasites. At 6 weeks post clearance, mice were challenged with a 2‘ *Hpb* inoculation and controls included naïve mice orally gavaged with PBS. At day 4 post 2‘ inoculation, small intestines were collected. 4uM frozen swiss-roll sections stained with AF488-F4/80 and AF647-CD206, Hoechst 33342 (a,b). Villin^Cre^-A_2B_AR^fl/fl^ and Villin^Cre^ mice were given a 2‘ *Hpb* inoculation as just described. At day 4 post 2‘ inoculation, punch biopsies of granulomatous tissue n=10 (c-j) were analyzed by RT-PCR. Data shown are the mean and SEM from 6 mice/group (One-way ANOVA, ****p<0.0001, (* comparisons as in Fig.1). Experiments were repeated at least two times with similar results.

To further examine effects on the tissue-dwelling larval phase, Villin^Cre^-A_2B_AR^fl/fl^ and Villin^Cre^ control mice were inoculated orally with 200 L3 *Hpb* (1‘*Hpb*) and 14 days post infection, mice were administered the anti-helminthic drug pyrantel pamoate via oral gavage for 2 consecutive days, effectively removing all parasites from the gut, as described previously [14]. At six weeks post clearance, Villin^Cre^-A_2B_AR^fl/fl^ and Villin^Cre^ control mice were challenged with a secondary inoculation of 200 L3 (2‘ *Hpb*). At this same time point, control groups of naïve Villin^Cre^-A_2B_AR^fl/fl^ ad Villin^Cre^ control mice were given a primary (1‘) *Hpb* inoculation. Seven days later all groups were analyzed for small intestinal expression of type 2 cytokines and M2 macrophage markers. We observed a significant reduction in IL-4, IL-13, and IL-5 mRNA and decreased M2 macrophage markers (Chi3I3, Retnla, and Arg1) in Villin^Cre^-A_2B_AR^fl/fl^ (Supplementary Fig. 2), indicating that IEC A_2B_AR deficiency impairs the induced localized secondary type 2 immune response at the host: parasite interface and also reduces effective protective immunity during the tissue dwelling stage.

### Memory IL-4 producing CD4 T cells are sufficient to mediate protective immunity in naïve mice

Previous studies have shown that the memory type 2 immune response to *Hpb* is IL-4 dependent [25], a finding we confirmed (Supplementary Fig. 3a). Furthermore, adoptive transfer of in vivo primed CD4 T cells to naïve mice can drive accelerated resistance similar to an intact memory response [14]. To corroborate and extend these studies, WT and IL-4^−/−^ mice were infected with *Hpb* and treated with pyrantel pamoate at day 14 after inoculation. Six weeks after primary inoculation 5×10^6^ CD4^+^ T cells, collected from mesenteric lymph nodes and spleen, were transferred to naïve recipient mice. Two days later, recipients were inoculated with *Hpb* and 14 days later worm burden was assessed. WT and IL-4^−/−^ mice that received CD4^+^ T cells from primed WT mice had a significant decrease in worm burden at day 14, but WT and IL-4^−/−^ mice that received CD4 T cells from primed IL-4^−/−^ mice had an impairment in worm expulsion and exhibited high worm burden (Supplementary Fig. 3b). These studies indicate that IL-4 produced by memory T cells (but not non-T cells) is necessary and sufficient to mediate acquired resistance in naïve recipient mice. To assess whether the donor memory CD4^+^ T cells from WT mice localized to the granuloma in peripheral intestinal tissue, 4get/KN2 IL-4 reporter mice were primed with *Hpb*, treated with pyrantel pamoate and at 14 days transferred to naïve WT recipients. At day 7 after inoculation small intestine swiss roll cryosections were analyzed for hCD2 (red, IL-4 protein marker) and GFP (IL-4 mRNA). Transferred hCD2-expressing CD4^+^ T cells migrated to the granuloma and surrounded the worms, consistent with a functional peripheral T cell response and the observed decreased worm burden (Supplementary Fig. 3c,d).

**Figure 3:**
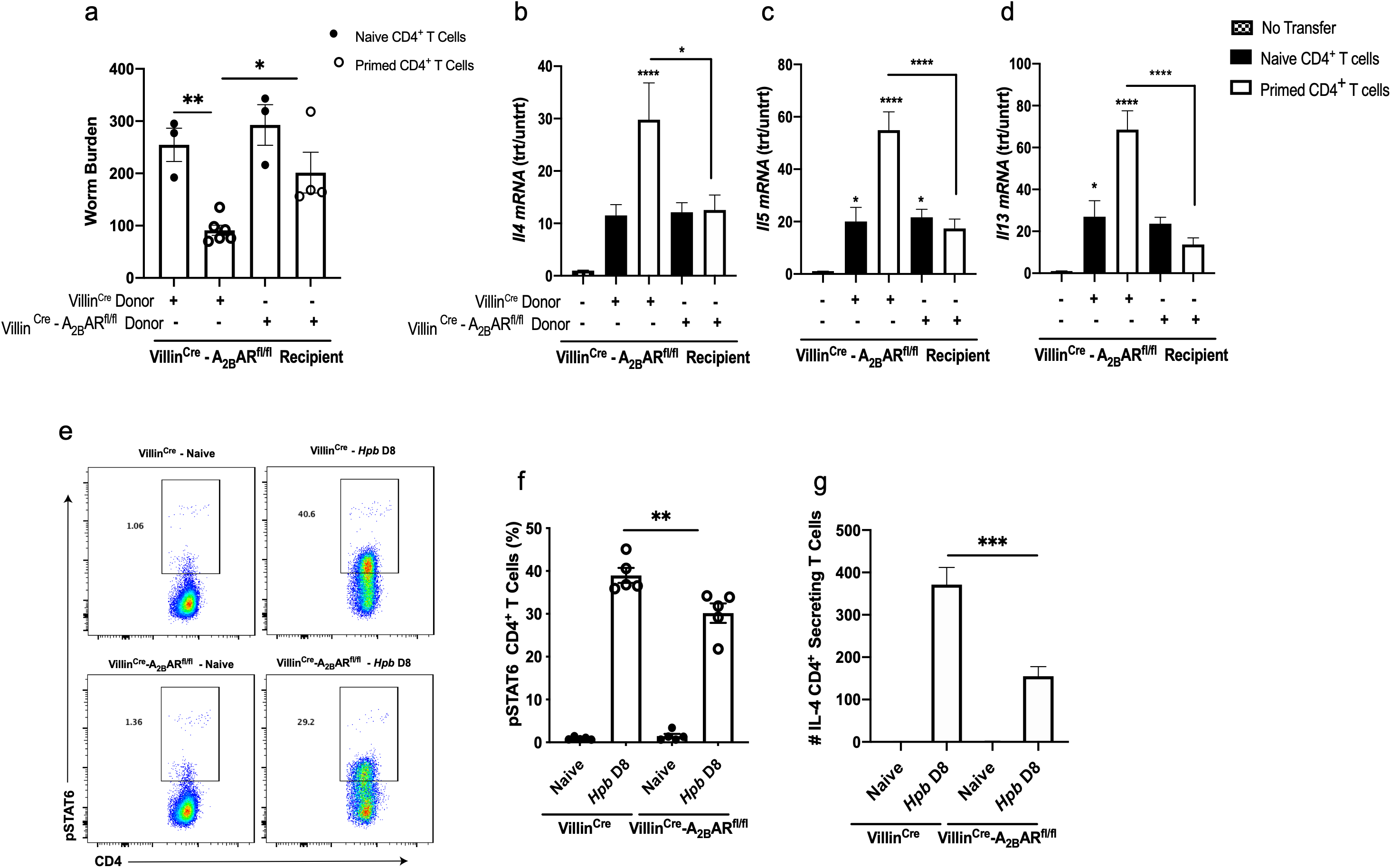
A_2B_AR deficiency in intestinal epithelial cells impaired memory CD4^+^ T cell function after primary *Hpb.* Villin^Cre^-A_2B_AR^fl/fl^ and corresponding control Villin^Cre^ mice were orally inoculated with 200 L3 *Hpb* and 14 days later, mice were treated with anti-helminthic drug, pyrantel pamoate to expulse parasites. At 6 weeks post clearance, CD4^+^ T cells were magnetically sorted from MLNs and spleens. 5×10^6^ CD4+ T cells from both treatment groups were retroorbital injected into naïve Villin^Cre^-A_2B_AR^fl/fl^ recipient mice. Controls included CD4+ T cells from naïve Villin^Cre^-A_2B_AR^fl/fl^ and Villin^Cre^ mice injected into naïve Villin^Cre^-A_2B_AR^fl/fl^ recipient mice. 2 days post transfer, mice were inoculated with 200 L3 *Hpb* larvae. 14 days post infection, worm burden was counted (a). Small intestine tissues (b-d) were analyzed by RT-PCR. Villin^Cre^-A_2B_AR^fl/fl^ mice and Villin^Cre^ corresponding controls were inoculated with 200 L3 *Hpb* larvae for 8 days. Mesenteric lymph node (MLNs) cell suspensions were used to measure expression of pSTAT6 in CD4^+^ T cells using flow cytometry. Representative flow cytometric analyses of CD4^+^ T cells expressing intracellular phosphorylated STAT6 (e) and percent of CD4^+^ T cells expressing pSTAT6 (f). CD4^+^ T cells from pooled mice were sorted and the frequency of IL-4 production was measured using ELISPOT assay (g). The number of CD4^+^ T cells are representative of quadruplicate wells (g). Data shown are the mean and SEM from 6 mice/group (One-way ANOVA ****p<0.0001, *comparisons as in Fig.1). Experiments were repeated at least 2 times with similar results.

### A_2B_AR deficiency in intestinal epithelial cells (IECs) impairs memory CD4^+^ T cell development but not subsequent activation

Our establishment of the above described T cell transfer model in the context of the *Hpb*-induced type 2 response provides a basis for further elucidating the role of A_2B_AR in the development of the type 2 memory response. Our findings indicated that T cells and the associated type 2 memory responses were impaired in *Hpb*-inoculated mice deficient in A_2B_AR on IECs. However, it remained unclear whether inhibition of the response was due to: 1) inhibition of initial memory T cell development during priming or 2) impaired subsequent activation of already formed memory T cells following secondary inoculation. To investigate effects of A_2B_AR deficiency in intestinal epithelial cells on memory T cell development, Villin^Cre^ and Villin^Cre^-A_2B_AR^fl/fl^ mice were inoculated with *Hpb* and 14 days later treated with pyrantel pamoate, which effectively removes all parasites. Six weeks after pyrantel pamoate treatment, 5×10^6^ CD4+ T cells were isolated from the MLNs and spleen from both treatment groups and transferred to naïve recipient Villin^Cre^-A_2B_AR^fl/fl^ mice. Two days after transfer recipient mice were inoculated with *Hpb* and 14 days after the primary *Hpb* inoculation, worm expulsion was assessed. As shown in Fig. 3a recipient mice with *Hpb*-primed CD4^+^ T cells from Villin^Cre^ WT had significant decreases in worm burden compared to mice receiving primed CD4^+^ T cells from Villin^Cre^-A_2B_AR^fl/fl^ mice. These findings suggest that the priming and development of CD4 memory Th2 cells in the context of helminth infection requires A_2B_AR signaling in intestinal epithelial cells. However, our findings that primed CD4+ T cells transferred to Villin^Cre^-A_2B_AR^fl/fl^ naïve mice could effectively mediate worm expulsion indicate that once CD4^+^ T memory cells have developed, A_2B_AR signaling on epithelial cells in not required for their subsequent activation and the associated development of a host protective response. Furthermore, gene expression of small intestine type 2 cytokines *Il-4*, *Il-5* and *Il-13*, were significantly increased in mice receiving primed CD4 T cells from Villin^Cre^ WT mice but not from CD4 T cells from Villin^Cre^-A_2B_AR^fl/fl^ mice (Fig. 3b-d). These studies indicate that the development of the memory T cell compartment during a primary response requires A_2B_AR signaling on IECs.

### A_2B_AR deficiency in IECs impaired IL-4R signaling in MLN CD4^+^ T cells after primary *Hpb* inoculation

Our finding that memory T cell development is impaired during the primary type 2 immune to *Hpb* raised the possibility that the overall type 2 response and associated development of CD4^+^ T cell effector cells were inhibited. CD4^+^ T cell STAT6 phosphorylation in the mesenteric lymph node is markedly elevated after *Hpb* infection consistent with increased localized bioavailability of IL-4. STAT6 phosphorylation was analyzed by flow cytometry on CD4 T cells in Villin^Cre^-A_2B_AR^fl/fl^ and Villin^Cre^ mice at day 8 after primary *Hpb* inoculation. As shown in Figure 3e,f significantly decreased STAT6 phosphorylation in CD4+ T cells was observed in *Hpb* infected Villin^Cre^-A_2B_AR^fl/fl^ compared to Villin^Cre^ mice. Furthermore, IL-4 protein secreted from sorted CD4^+^ T cells was also significantly reduced compared to WT infected Villin^Cre^ at day 8 after primary *Hpb* inoculation, as determined by ELISPOT (Fig. 3g). Similarly, CD4^+^ T cell pSTAT6 was also decreased as late as day 11 after both primary and secondary *Hpb* inoculation (Supplementary Fig. 3e,f). Taken together these studies indicate that A_2B_AR signaling in epithelial cells is also required for optimal Th2 effector cell development after primary inoculation.

### Epithelial A_2B_AR signaling is required during the initiation stages of Hpb primary infection

Our findings that A_2B_AR signaling on epithelial cells is required for the development of memory CD4^+^ T cells after primary inoculation but not for their subsequent activation after a secondary exposure suggested a critical role for A_2B_AR signaling in the development of memory T cells following an initial infection. *Hpb* can penetrate the epithelial barrier of the small intestine within hours after inoculation and induce type 2 cytokines and likely other as yet unidentified signaling pathways [26]. To address the role of epithelial cell A_2B_AR signaling in initiating this early response, Villin^Cre^-A_2B_AR^fl/fl^ mice and corresponding Villin^Cre^ controls were inoculated with *Hpb* and 24 hrs later small intestine was collected and RNA isolated for qPCR analyses. Consistent with previous studies, *Hpb*-inoculated Villin^Cre^ mice had a significant increase in the type 2 cytokines IL-4, IL-5, and IL-13, and the cytokine alarmin IL-33 in intestinal tissues. However, increases in these cytokines were largely blocked in *Hpb*-inoculated Villin^Cre^-A_2B_AR^fl/fl^ mice (Fig. 4a-d). Cytokines associated with type 1 immunity were also assessed for possible deviation of the response but increases in IFNγ and TNF were not markedly different in Villin^Cre^ and Villin^Cre^-A_2B_AR^fl/fl^ mice after *Hpb* inoculation (data not shown).

**Figure 4:**
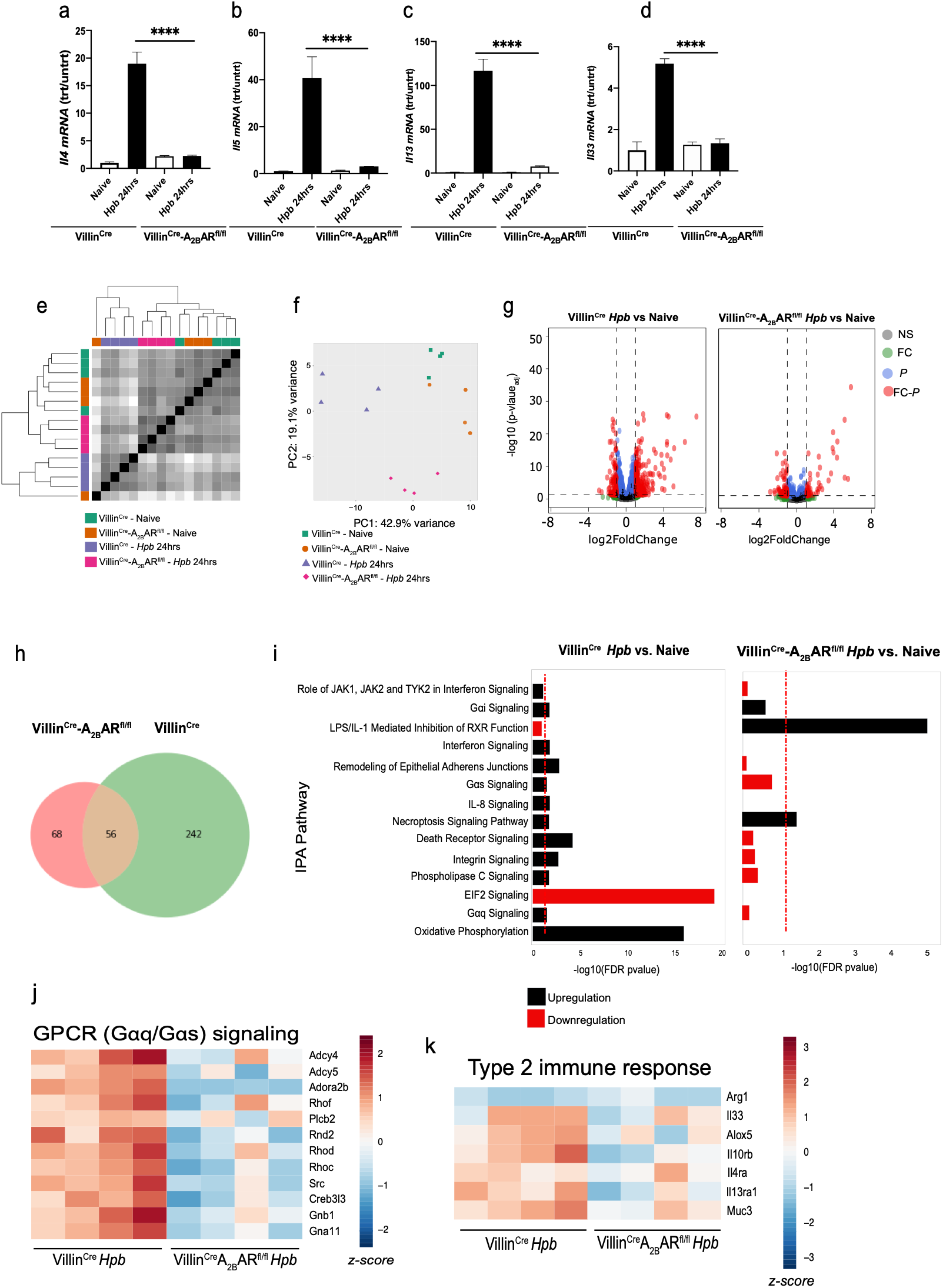
Epithelial A_2B_AR signaling is required during the initiation stages of Hpb infection. (a-d) Villin^Cre^-A_2B_AR^fl/fl^ mice and Villin^Cre^ corresponding controls were inoculated with 200 L3 *Hpb* larvae for 24 hours. Small intestinal tissue were analyzed by RT-PCR. Data shown are the mean and SEM from at least six individual mice per group (One-way ANOVA *p<0.01,* comparisons as in Fig.1). Experiments were repeated at least 2 times with similar results. (e-i) RNAseq analyses of intestinal epithelial cells (IECs) 24 hours after Hpb inoculation. Villin^Cre^-A_2B_AR^fl/fl^ and corresponding control Villin^Cre^ mice were orally inoculated with 200 L3 *Hpb* larvae for 24 hours, controls included naïve mice orally gavaged with PBS. Heat map representing Euclidean distance matrix (e) of total transcriptome of each sample, along with hierarchical clustering; principal component analysis (PCA) of total transcriptome of each sample; and volcano plots. Venn diagram depicting the number of upregulated and downregulated genes with a shared and unique expression from Villin^Cre^-A_2B_AR^fl/fl^ and corresponding controls Villin^Cre^ mice relative to Naïve (f). Enrichment Ingenuity Pathway Analysis (IPA) Pathway (g) Heat map representing GPCR signaling (h) and Type 2 immune response (i).

The profound decrease in activation of type 2 associated cytokine responses following A_2B_AR blockade on intestinal epithelial cells suggested that adenosine signaling on epithelial cells was likely critical at the very earliest stages of initiation of the type 2 immune response. To more globally assess changes in epithelial cell activation and associated stimulation of signaling pathways as a result of A_2B_AR deficiency, small intestinal epithelial cells (DAPI^−^ CD45^−^ EpCAM^+^) were sorted from Naive Villin^Cre^ and Villin^Cre^-A_2B_AR^fl/fl^ mice or at 24 hours after *Hpb* inoculation. Highly purified RNA was individually purified from 4 mice/treatment group and then subjected to RNAseq analyses. Euclidean distance matrix along with hierarchical clustering (Fig. 4e) and Principal component analysis (PCA) (Fig. 4f) of total transcriptomes from each sample, showed similar gene expression profiles of both naïve Villin^Cre^ and Villin^Cre^-A_2B_AR^fl/fl^ strains. However, *Hpb*-inoculated Villin^Cre^ and Villin^cre^-A_2B_AR^fl/fl^ mice showed distinct gene expression profile. Volcano plots (Fig. 4g) showed marked reduction in upregulated gene expression in Villin^Cre^A_2B_AR^FlFl^ compared to Villin^Cre^ after *Hpb* inoculation. Venn diagram analyses (Fig. 4h) depicting the number of upregulated and downregulated genes with a shared and unique expression from Villin^Cre^-A_2B_AR^fl/fl^ and corresponding controls Villin^Cre^ mice relative to naïve mice identified 242 genes that were differently expressed in WT mice and 60% of those upregulated while 68 genes were differently expressed in Villin^Cre^-A_2B_AR^fl/fl^ mice with 19% upregulated genes (Fig. 4h). Enrichment Ingenuity Pathway Analysis (IPA) (Fig. 4i) was used to identify upregulated signaling pathways. Oxidative phosphorylation, and Gs and Gq protein and phospholipase C (PLC) signaling were the most reduced in *Hpb*-inoculated Villin^Cre^-A_2B_AR^fl/fl^ compared to *Hpb*-inoculated VillinCre mice indicating that the Gs, Gq, and PLC pathways may mediate the A_2B_AR induction of the IEC response to *Hpb* (Fig. 4j). Furthermore, we identified type 2 immune response genes significantly reduced in infected Villin^Cre^-A_2B_AR^fl/fl^ compared to infected VillinCre mice (Fig. 4k). These findings thus indicate that after *Hpb* inoculation, A_2B_AR signaling is required for effective activation of epithelial cells essential for production of factors critical for optimal type 2 immune responses required for the development of an effective CD4 dependent memory response.

### Adenosine metabolized from extracellular ATP promotes type 2 immune response through triggering epithelial A_2B_AR signaling

Our findings indicate that shortly after *Hpb* infection epithelial cell activation is compromised when their expression of A_2B_AR is blocked. To examine the source of adenosine at this early stage of the response, we hypothesized that extracellular ATP may play a critical role. To determine the timing of ATP release after *Hpb* inoculation, we utilized novel transgenic ATP reporter mice that constitutively and ubiquitously express pmeLUC on the surface of all cells. Mice were inoculated with PBS or 200L3 *Hpb* larvae and imaged 24 hours post inoculation. 10 minutes before imaging, mice were injected intraperitoneally with D-luciferin (75 mg/kg). Our studies demonstrate ATP release within 24 hours post *Hpb* inoculation (Fig. 5a,b). Extracellular ATP can be immediately degraded into extracellular adenosine by ectonucleotidases CD39 and CD73 [27–32]. We have previously reported that intestinal epithelial lymphocytes increase surface expression of CD39 and CD73 within 24 hours of *Hpb* inoculation [9]. To examine whether CD39 played a significant role in the host protective memory response to *Hpb*, mice genetically deficient in CD39 and C57BL/6 wild-type controls were infected with *Hpb.* Specifically, mice were inoculated orally with 200 L3 *Hpb* (1‘*Hpb*) and 14 days post infection, mice were administered the anti-helminthic drug pyrantel pamoate via oral gavage for 2 consecutive days, effectively removing all parasites from the gut, as described previously [14]. At six weeks post clearance, CD39^−/−^ and C57BL/6 wild-type control mice were challenged with a secondary inoculation of 200 L3 (2‘ *Hpb*). At this same time point, control groups of naïve CD39^−/−^ and C57BL/6 wild-type controls mice were given a primary (1‘) *Hpb* inoculation. Fourteen days later all groups were analyzed for worm burden and egg counts. C57BL/6 wild-type mice had reduced worm burden consistent with an effective Th2 memory response, while CD39^−/−^ mice had significant increases in worm burden, indicating that the Th2 memory response was compromised (Fig.e 5c,d). To examine whether CD39 played a role in the initiation of type 2 immune response, CD39^−/−^ mice and controls were inoculated with *Hpb* and 24 hours later small intestine was collected and RNA isolated for qPCR analyses. *Hpb*-inoculated control mice had a significant increase in the type 2 cytokines and the cytokine alarmin IL-33 in intestinal tissues, and they were significantly impaired in CD39^−/−^ mice (Fig. 5e-h). CD39^−/−^ mice and C57BL/6 WT control mice were then infected with *Hpb* for 8 days. CD4^+^, pSTAT6^+^ T cells from MLNs of CD39^−/−^ mice were significantly decreased, as well as MHC II expression on B cells (Fig. 5i-j). Mice lacking CD39 also showed an impaired type 2 immunity to *Hpb* in the MLN (data not shown) small intestine (Fig. 5k-m), and granulomas (data not shown), as gene expression of type 2 cytokines and M2 macrophage markers were significantly decreased compared to WT mice. These results indicate that eATP is metabolized to adenosine, which then acts as an essential endogenous danger signal.

**Figure 5.**
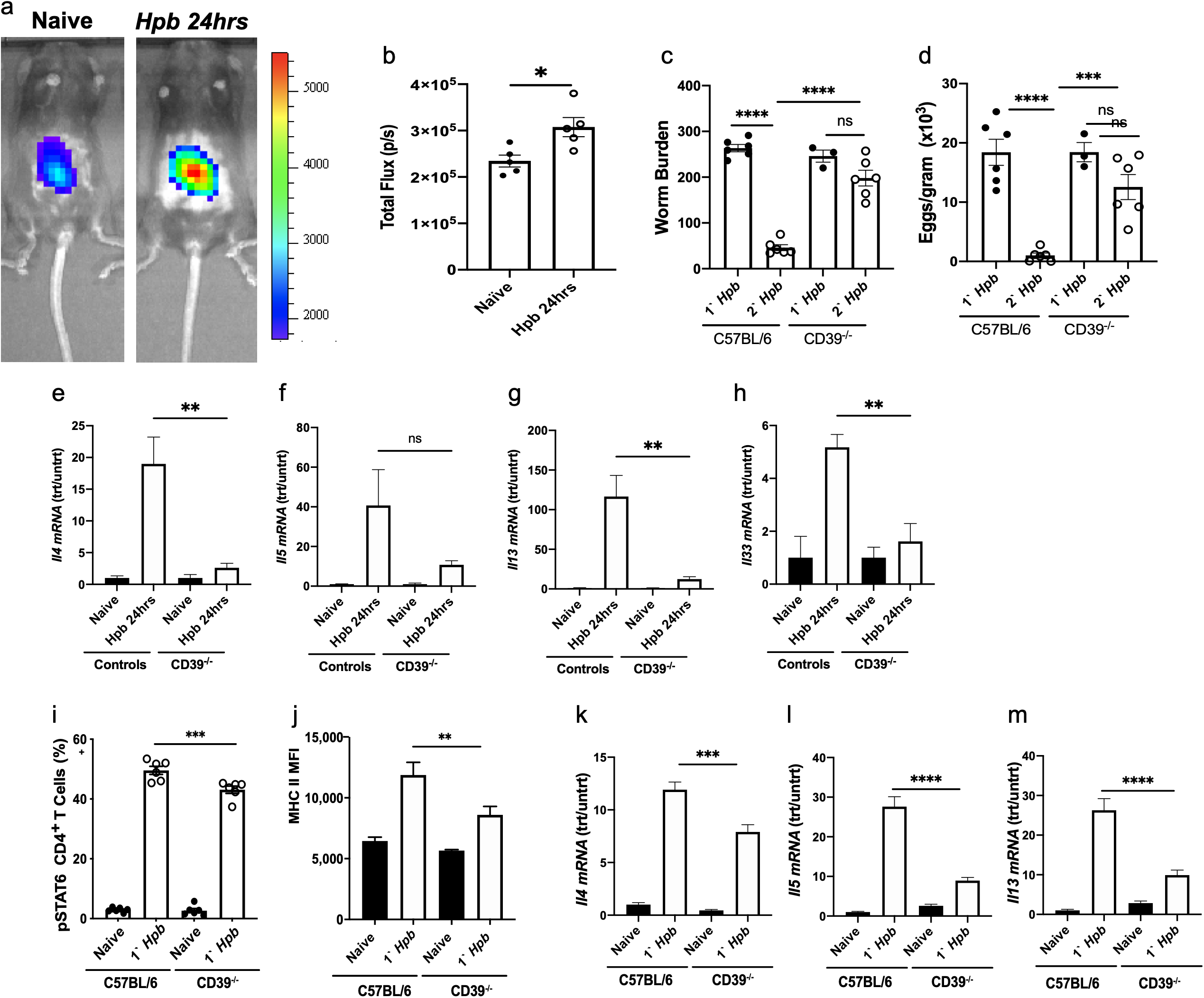
Adenosine metabolized from extracellular ATP promotes type 2 immune response through triggering A_2B_AR signaling on epithelial cells. pme-Luc mice were inoculated with 200 L3 *Hpb* larvae, control group included mice orally gavaged with PBS. 24 hours post gavage mice were imaged using Xenogen IVIS-200 System for detection of ATP (a), measured total flux (b). CD39^−/−^ mice and corresponding C57BL/6 controls were orally inoculated with 200 L3 *Hpb* larvae, 14 days later, mice were treated with anti-helminthic drug, pyrantel pamoate for 2 constitutive days to expulse the parasites. At 6 weeks post clearance, mice were challenged with a secondary (2‘) *Hpb* inoculation and controls included mice given primary (1‘) *Hpb* inoculation. Resistance was assessed at day 14, luminal worm burden (c) and eggs were enumerated (d). CD39^−/−^ mice and corresponding controls were inoculated with 200 L3 *Hpb* larvae for 24 hours. Small intestinal tissue was analyzed by RT-PCR (e-h). CD39^−/−^ mice and corresponding C57BL/6 controls were infected with 200 L3 *Hpb* larvae for 8 days. Mesenteric lymph nodes (MLNs) cell suspensions were used to measure expression of intracellular pSTAT6 in CD4^+^ T cells (i), and mean MFI of MHC-II+ expressing B cells (j) using flow cytometry. Small intestine (k-m) were analyzed by RT-PCR. Data shown are the mean and SEM from 4-5 mice/group (One-way ANOVA ****p<0.0001, * comparisons as in Fig.1). Experiments were repeated at least 2 times with similar results.

## Discussion

Our results indicate a significant role for adenosine specifically binding A_2B_AR on epithelial cells in driving a host protective memory immune response to the intestinal nematode, *Hpb*. We further show that A_2B_AR signaling is required for the development of memory CD4 T cells but their subsequent activation during a memory response is independent of this signaling pathway. During the primary response, A_2B_AR signaling augments the development of effector T cells and markedly promotes initial activation of innate immunity in the intestinal tissue shortly after infection.

It is likely that the early localized increased eATP observed within 24 hours after *Hpb* inoculation is a result of tissue damage associated with the parasites crossing the epithelial barrier. Previous studies have shown that P2X7 receptor binding of eATP on mast cells can drive their production of IL-33, and that in WT mice this is required for optimal host protective effects [6]. Our studies now show that eATP catabolized to adenosine plays an essential role in driving the initial innate type 2 immune response, further upregulating IL-33 production. Tuft cells have also been shown to play an important role in driving the type 2 response to helminths [3, 33–36]. During the early stages of infection tuft cells initially sense factors secreted by the helminth in the intestinal lumen, resulting in production of cysteine leukotrienes, which together with constitutive IL-25 initiate a tuft cell-ILC-2 circuit [3]. As the parasite actually invades the intestinal epithelium, tissue damage triggers the release and production of DAMPs, in particular eATP and adenosine respectively. Trefoil factors (Tff2 and Tff3) are also released after helminth infection acting as DAMPs to further promote IL-33 production [7, 8]. This combination then drives IL-33 production, which is essential for the maximal development of the type 2 immune response [37]. Taken together, these results suggest that multiple successive signals fine tune the upregulation of the type 2 immune response, thereby better controlling unnecessary inflammation-associated tissue damage.

As A_2B_AR signaling has previously been shown to promote M2 activation in vitro [10, 20, 21], we initially infected LysM^Cre^A_2B_AR^fl/fl^ mice with *Hpb* to specifically delete A_2B_AR in myeloid cells. Our findings that this had little effect on the immune response to *Hpb* indicated that other cell types were likely involved. We now demonstrate a critical role for epithelial cell A_2B_AR signaling in driving the primary and memory host protective response to this enteric helminth. Epithelial cells are likely of particular importance at barrier surfaces in triggering type 2 responses, allowing them to serve as sentinel cells capable of rapidly inducing innate immunity following sensing of helminth associated signals or as a result of direct cellular stress or damage. In contrast, deletion of A_2B_AR on myeloid cells blocked the development of the peritoneal type 2 immune response to sterile microparticles. This is consistent with previous studies showing an essential role for macrophages in producing IL-33 in this type 2 sterile inflammation model [23] and further demonstrates that in the context of this peritoneal model A_2B_AR signaling is now required on macrophage populations. It is likely that cellular stress/damage was a critical factor in stimulating the type 2 immune response following infection with multicellular parasites or inoculation with sterile microparticles. Thus, eATP may be released and catabolized to adenosine which then functions to initiate type 2 immunity. These findings indicate the importance of the specific tissue microenvironment in influencing the immune cell populations that actually trigger type 2 immune responses, with epithelial cells being critical at barrier surfaces and macrophages being essential in peritoneum. However, our data also demonstrate the general significance of A_2B_AR signaling in driving this type of response even though the tissue microenvironment and essential type 2 response-inducing cell types are distinct. Type 2 responses have now been shown to arise under a variety of different disease conditions including: cancer [38–41], microbial infections [42, 43], and traumatic injury [44–46]. It will be of interest in future studies to examine the role of A_2B_AR signaling in driving type 2 immunity in these very different responses and examining whether A_2B_AR signaling on epithelial cells, macrophages, or potentially other cell types is involved.

The ability of *Hpb* to stimulate a potent type 2 immune response during both the larval tissue dwelling phase and the subsequent adult luminal phase provides an opportunity to interrogate a broad spectrum of elicited immune responses. Also drug clearance after primary chronic infection eliminates all worms from the host and subsequent secondary inoculation results in a potent CD4^+^ T cell-dependent memory type 2 response, which includes resistance mechanisms activated during both the tissue-dwelling and the luminal phase, culminating in expulsion of worms by day 14 after secondary inoculation [14, 47]. Our analysis of the characteristic type 2 granuloma that develops at the host parasite interface following secondary inoculation showed profound decreases in M2 macrophages in mice lacking A_2B_AR signaling in epithelial cells. Previous studies have shown a critical role for M2 macrophages in impairing larval parasite development and enhancing subsequent worm expulsion [14]. As such, our findings of impaired granulomatous development in the tissue dwelling stage are consistent with the increased worm burden observed during the memory response in the *Hpb* inoculated Villin^Cre^A_2B_AR^fl/fl^ mice. As the parasite lifecycle includes only a short 8-day tissue-dwelling window, when macrophages and other cell types can potentially damage the developing larvae, the rapid activation of memory T cells that drive granuloma development is critical in mediating the accelerated resistance observed after secondary inoculation[48, 49]. Our adoptive transfer experiments revealed that epithelial A_2B_AR signaling is not required for activation of memory T cells after secondary inoculation. Previous studies have shown that memory T cell activation is less dependent on extrinsic signals, such as costimulatory molecules, than the development of memory T cells during the primary response [50]. Indeed, our findings showed that the development of memory T cells that could mediate accelerated parasite resistance after secondary exposure was blocked in the VillinCreA_2B_AR^fl/fl^ mice. Thus, A_2B_AR epithelial cell signaling is required for the development of memory Th2 cells following initial infection with helminth parasites.

Our observation, using pme-Luc reporter mice, that eATP is markedly increased at 24 hours after Hpb inoculation raised the possibility that the eATP may be an important source of adenosine, although it was also possible that intracellular adenosine secreted through the nucleoside transporter [51, 52] could be the primary source. eATP is metabolized to adenosine through CD39 and CD73. We had previously shown that CD39 and CD73 were upregulated shortly after *Hpb* inoculation, which generally correlates with increased release and degradation of ATP [9]. Our findings now demonstrate that accelerated resistance is blocked following secondary *Hpb* inoculation of CD39^−/−^ mice. Furthermore, immune cell activation and cytokine expression was also blocked following CD39 blockade, indicating that eATP is a critical source of the adenosine essential in driving the Hpb induced type 2 immune response. RNAseq analysis of IECs after *Hpb* infection showed that A_2B_AR signaling on epithelial cells triggered Gs and Gq proteins and phospholipase C signaling indicating that these pathways mediate A_2B_AR induction of the IEC response to *Hpb*. Furthermore, the blockade of increased IEC oxidative phosphorylation in *Hpb* inoculated Villin^Cre^A_2B_AR^fl/fl^ mice, characteristic of activated M2 macrophages, and the blockade of various type 2 response markers, and also IL-33, indicated that differential activation of epithelial cells in the context of type 2 immunity required adenosine interactions with IEC A_2B_ARs.

In summary our studies indicate that adenosine, derived from eATP, triggers IEC A_2B_AR signaling resulting in differential activation of epithelial cells. This activation, including IEC IL-33 upregulation, is required for the development of type 2 innate immunity and its absence impairs the development of memory Th2 cells, essential for acquired resistance.

## Acknowledgements

National Institutes of Health (NIH) grants R01DK113790 (W.C.G. and G.H.) and T32AI125185 to D.W.E.

## Disclaimer Statement

G.H. owns stock in Purine Pharmaceuticals, Inc. and has patents related to purinergic signaling in sepsis.

## Methods

### Mice

8-12 week old C57BL/6 WT mice were purchased from The Jackson Laboratory (Bar Harbor, ME). We generated and have described the Villin^Cre^-A_2B_AR^fl/fl^ mice and LysM^Cre^-A_2B_AR^fl/fl^ mice (C57BL/6)[53], pme-Luc [54], and CD39^−/−^[55], as well as Villin^Cre^ [53] and LysM^Cre^ [54] mice previously. All mice were maintained in a specific pathogen-free, virus Ab-free facility during the experiments. Female and male healthy mice were selected for treatment groups from purchased or bred colonies, without using specific randomization methods or specific blinding methods. The studies have been reviewed and approved by the Institutional Animal Care and Use Committee at Rutgers-the State University of New Jersey. The experiments herein were conducted according to the principles set forth in the Guide for the Care and Use of Laboratory Animals, Institute of Animal Resources, National Research Council, Department of Health, Education and Welfare (US National Institutes of Health).

### Parasite inoculations

*H. polygyrus bakeri (Hpb)* L3 larvae were isolated from cultures using a modified Baermann apparatus and maintained in PBS at 4°C. To assess memory responses, mice were inoculated with by oral gavage (per os, p.o.) with 200 infective L3 larvae to establish a primary chronic infection. At day 14, mice were treated using an anti-helminthic drug (pyrantel pamoate 1mg/mouse) to expulse the parasites. 6 weeks post clearance, 200 L3 larvae is administered again as a challenge infection and naïve mice will be infected for a primary infection for 14 days. Worm burden, fecundity determined from fecal contents at day 14 post primary and challenge infections as described previously [14].

### Total ATP Assay and Protein Measurement

Total ATP Assay Five female worms were isolated after *H. polygyrus bakeri* inoculation of (Villin^Cre^-A_2B_AR^fl/fl^) and (LysM^Cre^-A_2B_AR^fl/fl^) mice, incubated in 100μl PBS with 100μl Cell-Glo reagents (Promega, Madison), and ground with a motorized pestle. After centrifugation at 5,300 rpm for 5 min, 100μl supernatant was transferred to wells of 96-well plate and luminescence measured in SpectraMax i3X cytometer. Controls included the following: PBS alone, heat killed worms; worms incubated without Cell-Glo reagent.

### Cytokine gene expression by RT-PCR

For qPCR, RNA was extracted from small intestine, MLN, granuloma punches and reverse transcribed to cDNA. qPCR was performed with Taqman (Applied Biosystems) kits and the Applied Biosystems 7500 Real-Time PCR System. All data were normalized to 18S ribosomal RNA, and the quantification of differences between treatment groups was calculated treated/untreated. Gene expression is presented as the fold increase over naïve controls.

### Flow cytometry

Flow cytometry MLN cell suspensions were collected and prepared from Hp1‘ and 2‘ inoculated WT and Villin^Cre^-A_2B_AR^fl/fl^ and LysM^Cre^-A_2B_AR^fl/fl^ mice; washed; blocked with Fc Block; and stained with anti-CD4-APC (RM4-5,), anti-pSTAT6-PE (pY641), anti-B220-PE(AR3-6B2), anti-CD3e-BV421 and anti-MHCII-percp/cy5.5 (IA/IE). Phosphorylation of STAT6 at tyrosine 641 was detected by intracellular staining with PE-conjugated anti-phospho-STAT6 using PhosFlow Fix Buffer I and Perm Buffer III reagents. Cells were acquired on Fortessa X-20 Flow cytometer (BD Biosciences, San Jose), and analyzed using FlowJo software.

### Adoptive Transfers

CD4^+^ T cells from MLNs of *Hpb* cured Villin^Cre^-A_2B_AR^fl/fl^ and corresponding controls Villin^Cre^ infected mice were magnetically sorted from MLN and spleen single cell suspensions using mouse CD4 MicroBeads (Miltenyi Biotec, Inc.) sorted on positive selection columns. 5 × 10^6^ cells retro-orbitally injected into sex- and age-matched Villin^Cre^-A_2B_AR^fl/fl^ recipients, which were infected primary *Hpb* inoculation 2 days after transfer.

### CD4+ T cells sorting and ELISPOT

Single cell suspensions from mesenteric lymph nodes IL-4 ELISPOT were performed using mouse IL-4 ELISPOT pair antibodies (BD Bioscience) 200,000 cells/well were cultured on multiScreen IP filter plate (Millipore) for 24 hours. Plates were scanned using C.T.L C6 Flourospot and Number of IL-4 secreting CD4 T cells were counted on counted on Immunospot software.

### In vivo ATP measurement

Transgenic pme-LUC mice were orally inoculated with 200 L3 *Hpb* larvae or PBS, 24 hours later, 75 mg/kg VivoGlo luciferin (Progema) was injected *i.p.,* 10 minutes later, whole body luminometry performed using IVIS 200 preclinical *in vivo* imaging system.

### Immunofluorescent staining

Fresh swiss-rolls of duodenum of the small intestine were O.C.T imbedded and flash frozen in 2-Methylbutane/liquid nitrogen and stored in −80°C. 4-5 μm sections were fixed in acetone for 8 minutes, blocked with 1% rat serum/1% Fc Block and stained with F4/80-AF488, CD206-AF647 (5μg/ml), Hoechst 33342 and sealed with ProLong Gold Antifade (Invitrogen). Images were taken using a Leica DM6000B fluorescent microscope, Orca Flash 4.0 mounted digital camera (Hamamatsu Photonics K.K., Japan) and LAS Advanced Fluorescence software (Leica Microsystems, Buffalo Grove, IL). Fluorescent channels were photographed separately and then merged. Exposure times and fluorescence intensities were normalized to appropriate control images.

### Bulk RNA sequencing

Intestinal epithelial cells from Villin^Cre^-A_2B_AR^fl/fl^, Villin^Cre^ *Hpb* infected for 24hrs and Naïve controls mice were sort-purified by gating on DAPI-, CD45-, EpCAM+ IECs. 4 biological replicates from each group was sorted purified individually. The purity of cell populations was 98% or greater. RNA was extracted using the RNeasy Mini Kit (QIAGEN). Illumina compatible NEB Next kit was used for library prep and sequenced using an Illumina NovaSeq6000 using NovaSeq SP kit with PE 2×100 configuration. Raw transcriptome reads were assessed for quality control (FASTQC v0.11.8) and trimmed for quality and adapter contaminant (cutadapt v 2.5). Trimmed reads were aligned to the mouse genome (GRCm38) using STAR (v2.6.1), followed by transcript abundance calculation and hit count extraction with StringTie (v2.0) and featureCounts (v1.6.4) respectively. Hit count normalization and differential gene expression group cross-comparisons were performed using DESeq2 (v1.26.0). Significant differentially expressed gene thresholds were set at FDR adjusted p <0.05. Pathway enrichment was performed using Ingenuity Pathway Analysis (Qiagen).

### Quantification and Statistical Analysis

Data were analyzed using the statistical software program Prism (GraphPad Software, Inc., La Jolla, CA) and are reported as means (±SEM). Differences between two groups were assessed by Student’s t test, differences among multiple groups were assessed by one-way ANOVA and individual two-way comparisons were analyzed using Tukey’s multiple comparison test. Differences of *p* < 0.05 were considered statistically significant.

---

**Supplementary Figure 1:**
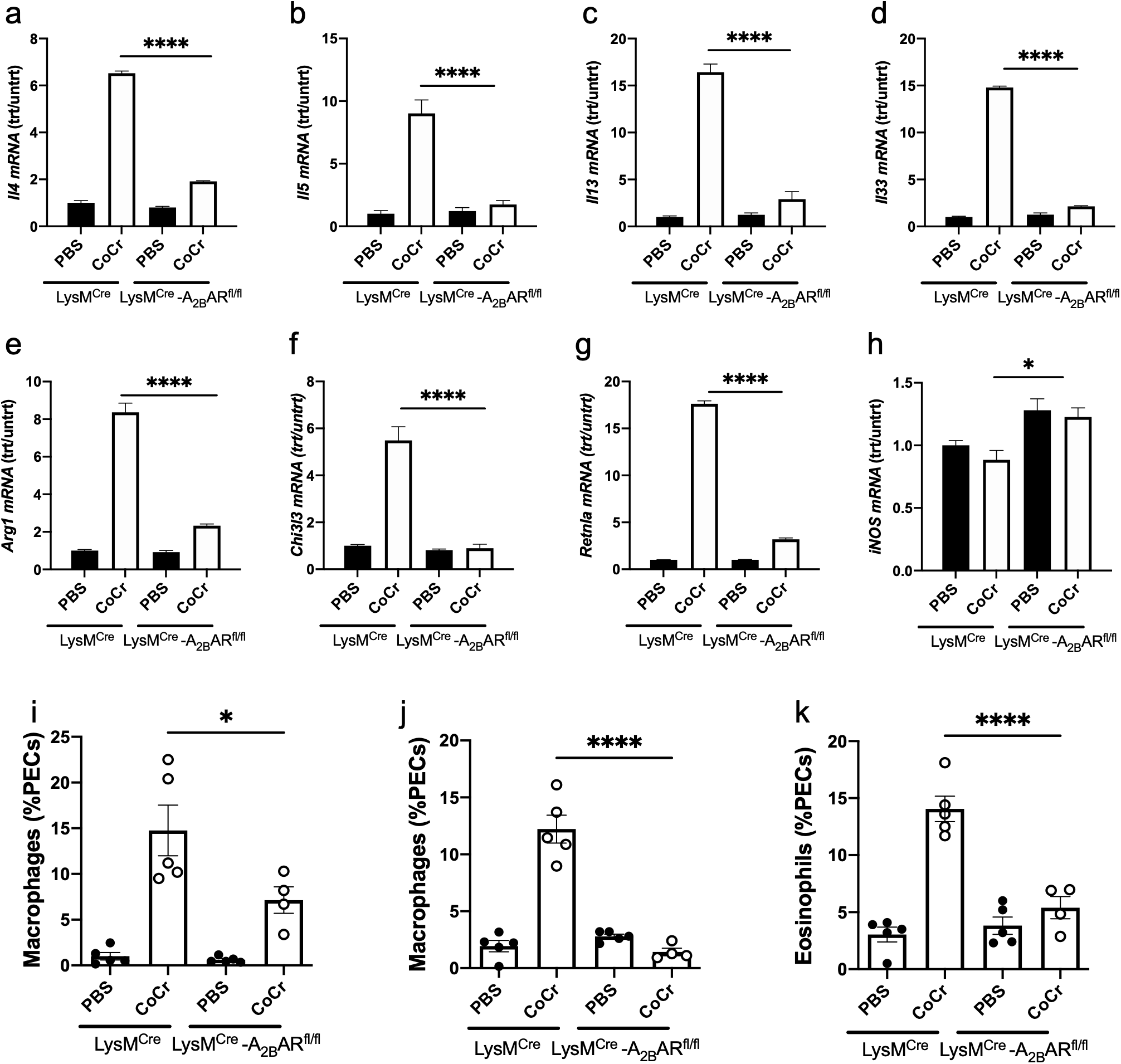
A_2B_AR signaling on myeloid cells is required for peritoneal type 2 immune response to sterile inert microparticles (MPs). LysM^cre^ -A_2B_AR^fl/fl^ mice and corresponding control LysM^Cre^ mice were injected *i.p.* with PBS vehicle or CoCr for 48 hours. Peritoneal cells were analyzed by RT- PCR (a-h) and assessed by flow cytometry for neutrophils (CD11b+, Ly6G+)(o), eosinophils (c-Kit-, Siglec-F+) (j), and M2 macrophages (F4/80+, CD206+) (k) as percentage (mean and se) of PECs. The mean and SE is shown for 4-5 mice/treatment group. One-way ANOVA ****p<0.0001, * comparisons as in Fig.1). Experiments were repeated at least 2 times with similar results.

**Supplementary Figure 2:**
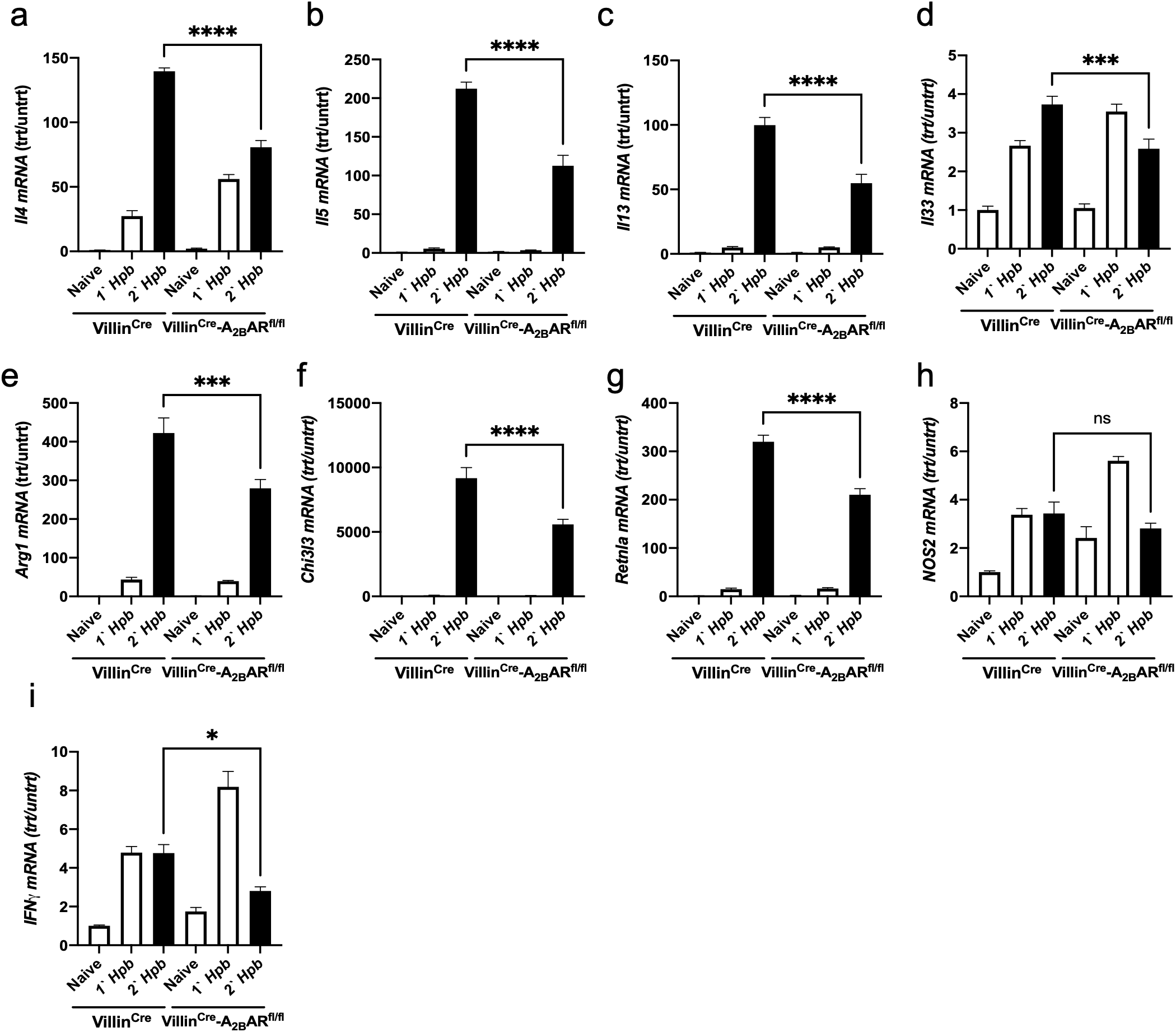
A_2B_AR IEC deficiency inhibits upregulated Th2 cytokine genes. Villin^Cre^-A_2B_AR^fl/fl^ and corresponding controls Villin^Cre^ mice were orally inoculated with 200 L3 *Hpb* larvae, 14 days later, mice were treated with anti-helminthic drug, pyrantel pamoate, to expulse the parasites. At 6 weeks post clearance, mice were challenged with a secondary (2‘) *Hpb* inoculation and controls included mice given primary (1‘) *Hpb* inoculation and naïve mice orally gavaged with PBS at day 7 post 1‘ and 2‘ inoculation. Small intestine mRNA was analyzed by qRT-PCR (a-i). Data shown are the mean and SEM from at least six individual mice per group (One-way ANOVA *p<0.01, *comparisons as in Fig.1). Experiments were repeated at least 2 times with similar results.

**Supplementary Figure 3:**
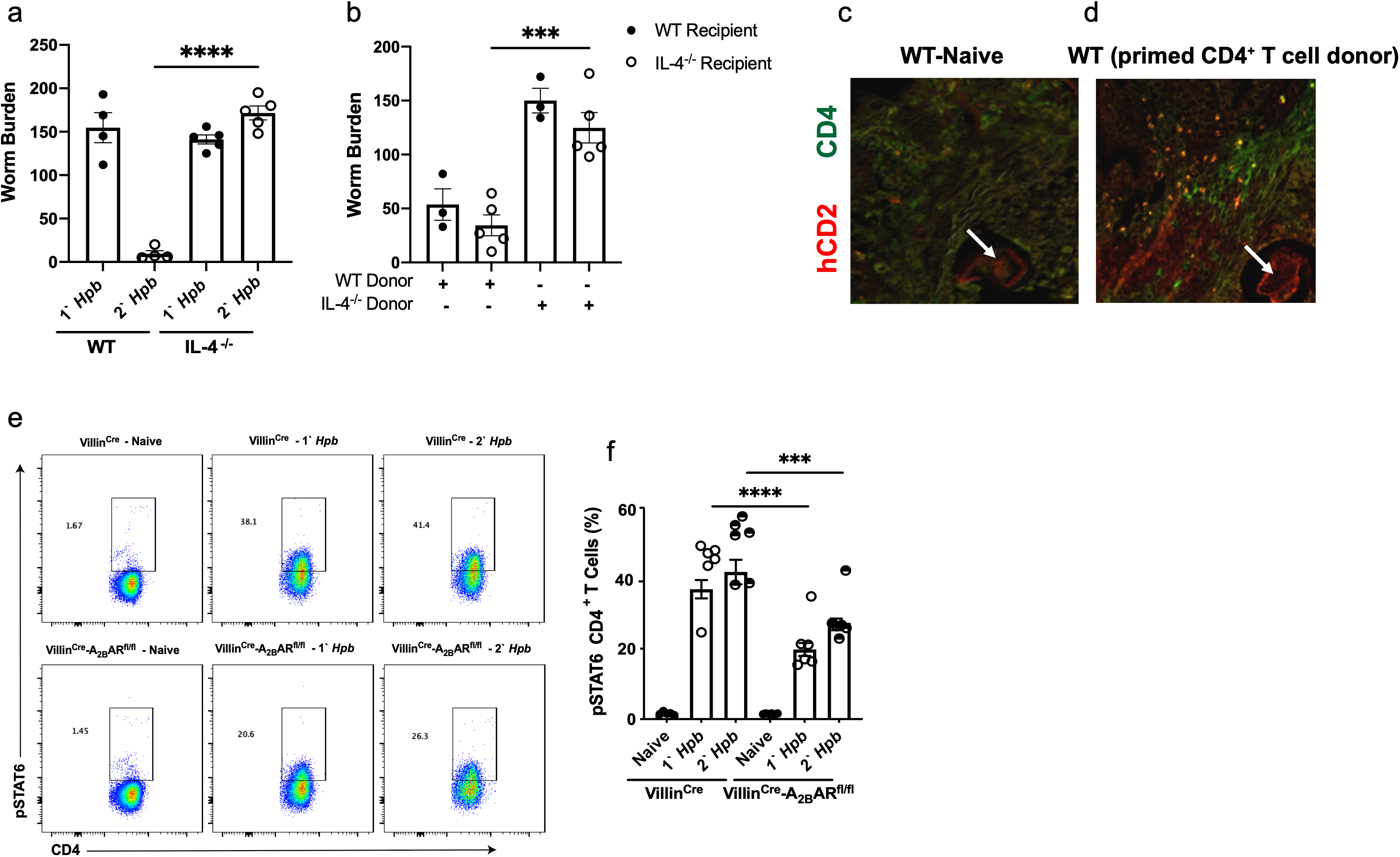
Transferred memory T cells can mediate acquired resistance in naïve recipients and A_2B_AR deficiency in intestinal epithelial cells impaired CD4^+^ T cell cytokine expression and responsiveness after challenge *Hpb* inoculation. IL-4^−/−^ andcontrol BL/6 mice were orally inoculated with 200 L3 *Hpb* larvae and 14 days later, mice were treated with anti-helminthic drug, pyrantel pamoate to expulse the parasites. At 6 weeks post clearance, mice were challenged with a secondary (2‘) *Hpb* inoculation and controls included mice given primary (1‘) *Hpb* inoculation. Resistance was assessed at day 14, luminal worm burden was enumerated (a). IL-4^−/−^ and corresponding controls mice were orally inoculated with 200 L3 *Hpb* larvae, 14 days later, mice were treated with anti-helminthic drug, pyrantel pamoate for 2 constitutive days to expulse the parasites. At 6 weeks post clearance, CD4^+^ T cells were magnetically sorted from MLNs and spleens. 5×10^6^ CD4+ T cells from both treatment groups were retroorbital injected into naïve WT and Naïve IL-4^−/−^ recipient mice. 2 days post transfer, mice were inoculated with 200 L3 *Hpb* larvae. 14 days post infection, worm burden was counted (b). CD4+ T cells were isolated from Naïve or *Hpb*-primed 4get/KN2 mice, and transferred into WT or IL-4^−/−^ recipients. After two days, recipients were given *Hpb* inoculation for 7 days. Small intestines swiss roll sections were stained CD4 (green) and hCD2 (red). Data shown from 4 mice/group. Villin^Cre^-A_2B_AR^fl/fl^ and corresponding control Villin^Cre^ mice were orally inoculated with 200 L3 *Hpb* larvae, 14 days later, mice were treated with anti-helminthic drug, pyrantel pamoate to expulse the parasites. At 6 weeks post clearance, mice were challenged with a secondary (2‘) *Hpb* inoculation and controls included mice given primary (1‘) *Hpb* inoculation and naïve mice orally gavaged with PBS. At day 11 post 1‘ and 2‘ inoculations, MLN cell suspensions were used to measure expression of intracellular pSTAT6 in CD4+ T cells using flow cytometry. Representative flow cytometric analyses of CD4+ T cells expressing phosphorylated STAT6 (e) and percent of CD4^+^ T cells expressing pSTAT6 (f). Data shown for both experiments are the mean and SEM from at least six individual mice per group and are representative of at least two independent experiments (One-way ANOVA * p<0.01 Villin^Cre^-A_2B_AR^fl/fl^ and corresponding controls Villin^Cre^ mice were orally inoculated with 200 L3 *Hpb* larvae, 14 days later, mice were treated with anti-helminthic drug, pyrantel pamoate for 2 constitutive days to expulse the parasites. At 6 weeks post clearance, mice were challenged with a secondary (2‘) *Hpb* inoculation and controls included mice given primary (1‘) *Hpb* inoculation and naïve mice orally gavaged with PBS. At day 11 post 1‘ and 2‘ inoculations, MLN cell suspensions were used to measure expression of intracellular pSTAT6 in CD4+ T cells using flow cytometry. Representative flow charts of CD4+ T cells expressing phosphorylated STAT6 (e) and percent of CD4^+^ T cells expressing pSTAT6 (f). Data from both experiments show the mean and SEM from at least six individual mice per group and are representative of at least two independent experiments (One-way ANOVA *p<0.01, *comparisons as in Fig.1).

